# Age-Related Changes of Dopamine D1 and D2 Receptors Expression in Parvalbumin-Positive Cells of the Orbitofrontal and Prelimbic Cortices of Mice

**DOI:** 10.1101/2023.12.17.571962

**Authors:** Jihui Dong, Xiaoyan Wei, Ziran Huang, Jing Tian, Wen Zhang

**Author notes:** Contact Information: Dr. Wen Zhang National Institute on Drug Dependence, Peking University, 38 Xueyuan Road, Beijing, 100191, China.

## Abstract

Dopamine (DA) plays a pivotal role in reward processing, cognitive functions, and emotional regulation. The prefrontal cortex (PFC) is a critical brain region for these processes. Parvalbumin-positive (PV+) neurons are one of the major classes of inhibitory GABAergic neurons in the cortex, they modulate the activity of neighboring neurons, influencing various brain functions. While DA receptor expression exhibits age-related changes, the age-related changes of these receptors in PV+ neurons, especially in the PFC, remains unclear. To address this, we investigated the expression of DA D1 (D1R) and D2 (D2R) receptors in PV+ neurons within the orbitofrontal (OFC) and prelimbic (PrL) cortices at different postnatal ages (P28, P42, P56, and P365). We found that the expression of D1R and D2R in PV+ neurons showed both age- and region-related changes. PV+ neurons in the OFC expressed a higher abundance of D1 than those in the PrL, and those neurons in the OFC also showed higher co-expression of D1R and D2R than those in the PrL. In both the OFC and PrL, D1R in PV+ neurons increased from P28 and reached a plateau at P42, then receded to express at P365. Meanwhile, D2R did not show significant age-related changes in both regions. These results showed dopamine receptors in the prefrontal cortex exhibit age- and region-specific changes, which may contribute to the difference of these brain regions in reward-related brain functions.

## 1 Introduction

The prefrontal cortex (PFC) plays a central role in various cognitive processes, including working memory, decision-making, attention, emotion, and memory (Dias, Robbins, & Roberts, 1996; Fuster & Alexander, 1971; Gold & Shadlen, 2007; Goldman-Rakic, 1987; Ochsner & Gross, 2005). It is vulnerable to multiple neurological and psychological diseases, such as depression, schizophrenia, and addiction (Belmaker & Agam, 2008; Koob & Volkow, 2010; Lewis, Hashimoto, & Volk, 2005). The PFC of rodents can be subdivided into four primary subregions, extending from dorsal to ventral: cingulate, prelimbic (PrL), infralimbic (IL), and orbitofrontal (OFC) cortices (Brodmann, 1909; Caviness, 1975; J. E. Rose & C. N. Woolsey, 1948; Jerzy E. Rose & Clinton N. Woolsey, 1948; Uylings & Van Eden, 1990; Zilles, 2012). While both the PrL and OFC are involved in decision-making and emotional regulation (Hernandez et al., 2022; Mızrak, Bouffard, Libby, Boorman, & Ranganath, 2021; Zeeb, Baarendse, Vanderschuren, & Winstanley, 2015), they exhibit distinct functions. The PrL is more associated with executive functions (Baker & Ragozzino, 2014; Broschard, Kim, Love, Wasserman, & Freeman, 2021) and emotional regulation (Mears, Boutros, & Cromwell, 2009), whereas the OFC plays a particular role in the evaluation of rewards and punishments, guiding appropriate behavioral responses (Frontera et al., 2023; Lichtenberg et al., 2021). Dysfunction in the PrL has been implicated in disorders such as anxiety (Luo et al., 2023). In contrast, damage to the OFC can lead to changes in social behavior and decision-making (Bolla et al., 2003; du Plessis et al., 2018). The PFC undergoes age-related changes both in cognitive functions and brain substrates (Chao & Knight, 1997; West, 1996). The PFC showed structure changes with increased functional connections during development, in meantime the working memory also showed an increase toward puberty in human and primates (Bunge, Dudukovic, Thomason, Vaidya, & Gabrieli, 2002; Gathercole, Pickering, Ambridge, & Wearing, 2004; Zhou et al., 2014). And executive functions, such as working memory and prefrontal brain regions show disproportionately strong age-related declines compared with other cognitive functions (Salthouse, Atkinson, & Berish, 2003; Zhou et al., 2014).

In the cortex, the orchestrated activity of inhibition and excitation is critical for brain functions (Zhang et al., 2017; Zhang, Peterson, Beyer, Frankel, & Zhang, 2014). Recent studies have shown that disinhibition is implicated in several psychological disorders, such as schizophrenia (Kokkinou et al., 2021), depression (Kantrowitz et al., 2021) and attention-deficit/hyperactivity disorder (Niedermeyer, 2001). Cortical inhibition is mainly mediated by GABAergic interneurons, the major types of which are Parvalbumin-, Somatostatin-, and Vasoactive intestinal peptide-positive (PV+, SST+, and VIP+, respectively) interneurons. These neurons show diverse morphology, axon targeting location, and functional differences, which are manifested by their roles in circuit and brain functions, such as oscillation. Of these neurons, PV+ neurons show a consistent inhibitory effect on local excitatory pyramidal neurons (Zhang et al., 2016). Such a property makes PV+ neurons critical for cortical inhibition and functions, and studies have shown that changes in PV+ neuron activity or the synapse strength of PV+ to layer 5 excitatory neurons changed cortical output and brain functions (A. T. Lee et al., 2014; Sempere-Ferràndez, Martínez, & Geijo-Barrientos, 2019).

Both the PrL and OFC are crucial components of a vital brain circuit known as the reward pathway (Zhang, 2020). In the reward pathway, dopamine serves as a pivotal neurotransmitter. Dopamine is suggested as a reward signal in the brain for activities that controls animal actions, decisions, and choices, and acting as a reward signal that signals the discrepancy between the actual reward and its prediction (Farrell, Lak, & Saleem, 2022; J. Y. Lee et al., 2021; Seitz, Hoang, DiFazio, Blaisdell, & Sharpe, 2022). Dopamine receptors are G protein-coupled receptors, classified into two main types: D1-like receptors, including D1 and D5 subunits, and D2-like receptors, comprising D2-D4 subunits. These two types of receptors engage in distinct downstream signaling cascades within neurons. D1-like and D2-like receptors have opposing effects on adenylyl cyclase activity and cAMP concentration, as well as on phosphorylation of Dopamine- and cAMP-regulated neuronal phosphoprotein (DARPP-32) (Greengard, Allen, & Nairn, 1999; Hemmings & Greengard, 1986; Nishi, Snyder, & Greengard, 1997). By phosphorylation (facilitated by D1-like and inhibited by D2-like receptors), DARPP-32 inhibits the protein phosphatase PP-1, which modulates the activity of various voltage-gated and synaptic ion channels. For example, activation of D1-like receptors increased the intrinsic excitability of neurons, including PV+ neurons (Plateau, Baufreton, & Le Bon-Jego, 2023; Potts & Bekkers, 2022). Interestingly, in adolescent rats the modulation is exclusively D1- mediated, while in older animals a D2-mediated modulation is synergistic with the D1-mediated effect (Tseng & O’Donnell, 2007). For synaptic transmissions, dopamine enhances NMDA synaptic currents via D1 receptor and reduces via D2 receptor in the PFC (Banks et al., 2015; Chen & Yang, 2002), it also enhances GABAergic currents via D1-like receptors and reduces them via D2-like receptors in the striatum and PFC (Ji et al., 2009; Seamans, Gorelova, Durstewitz, & Yang, 2001).

Given the modulatory effects of D1-like and D2-like dopamine receptors on neuronal activity and the crucial role of PV+ interneurons in the circuit activities of cortical regions, it is essential to delineate the age-related changes of dopamine receptor expression profiles in PV+ neurons within the PrL and OFC to comprehend their contributions to brain functions. To address this question, we examine the expression patterns of dopamine D1 and D2 receptors in PV+ neurons in the OFC and PrL and compare age-related expression changes of these receptors in mice.

## 2 Materials and Methods

### 2.1 Animals

Male C57BL/6J mice (RRID: IMSR_JAX:000664) were used. Mice were maintained on a 12-hour light/dark cycle with food and water *ad libitum*. All experiments were performed in the dark cycle.

All procedures are in accordance with the National Institutes of Health *Guide for the Care and Use of Laboratory Animals* and have been approved by Peking University Animal Care and Use Committee.

### 2.2 Histology and immunofluorescence

For immunostaining, mice were anesthetized with isoflurane, then perfused with phosphate-buffer saline (PBS, pH 7.4) followed with 4% paraformaldehyde (PFA) in PBS. Brains were dissected and post-fixed with 4% PFA in PBS overnight at 4 °C, and 25-μm coronal sections were prepared with a vibratome. Immunostaining followed the standard protocols for free-floating sections. In brief, free-floating sections were incubated in blocking solution containing 4% normal donkey serum, 1% bovine serum albumin (BSA), and 0.3% Triton X-100 in PBS for 2 h at 23–25 °C. Sections were then treated with primary antibodies in blocking solution for 24 – 48 h at 4 °C, followed with secondary antibodies in blocking solution at 23–25 °C for 2 h with slow shaking.

Primary antibodies used were Goat Anti-Parvalbumin (1:2000, Swant, Cat# PVG-213, RRID: AB_2650496), Rat Anti-Dopamine D1 Receptor (1:200, Sigma, Cat# D2944, RRID: AB_1840787), Rabbit Anti-Dopamine D2 Receptor (1:250, Merck, Cat# AB5084P, RRID: AB_2094980).

Secondary antibodies used were Alexa Fluor 546 Anti-Goat (1:300, Thermo Fisher Scientific, Cat# A11056, RRID: AB_2534103), Alexa Fluor 488 Anti-Rat (1:300, Abcam, Cat# ab150153, RRID: AB_2737355), Alexa Fluor Plus 647 Anti-Rabbit (1:300, Thermo Fisher Scientific, Cat# A32795, RRID: AB_2762835).

### 2.3 Imaging

We acquired fluorescent images with a confocal microscope (Leica TCS-SP8 STED) using a 63× objective (NA 1.4) and a 16× objective (NA 0.5). The analysis was performed as previously described (Zhang et al., 2016). Briefly, a maximal projection of a 7 μm thick stack was analyzed with ImageJ (v1.53t, RRID:SCR_003070) based FIJI (RRID:SCR_002285) (Whitesell et al., 2021). The punta of D1 and D2 dopamine receptors of a minimum 2 pixels on parvalbumin-positive cell soma were analyzed with particle analysis of FIJI. The soma areas of PV+ cell, D1, and D2 dopamine receptors puncta size more than 5 times of the standard deviation were excluded from further analysis.

### 2.4 Statistical analysis

All statistical analyses and data plotting were performed with R (v4.2.2, RRID: SCR_001905), and the non-base attached packages for R were ggpubr (v0.6.0, RRID:SCR_021139), rstatix (v0.7.2, RRID:SCR_021240), tidyverse (v2.0.0, RRID:SCR_019186), and emmeans (v1.8.5, RRID:SCR_018734). For boxplots, whiskers denoted 1.5 * IQR from the hinges, which corresponded to the first and third quartiles of distribution. For multiple groups, one-way or two-way ANOVAs with *post hoc* Tukey’s test were used based on experiment design. n, sample number of cells; N, sample number of mice. *P* < 0.05 is considered statistically significant.

## 3 Results

### 3.1 The age-related changes of Parvalbumin-positive neurons in the orbitofrontal and prelimbic cortices

To examine the age-related expressions of dopamine D1 and D2 receptors (D1R and D2R, respectively) in parvalbumin-positive neurons in the orbitofrontal (OFC) and prelimbic (PrL) regions of the prefrontal cortex (PFC), we first examined parvalbumin expression in these two brain regions at post-natal days 14, 28, 42, 56, and 365 (P14, P28, P42, P56, and P365, respectively; **Figure 1**). We found that at P14, parvalbumin was barely expressed in the PrL, while the OFC showed strong parvalbumin expression. The cell density at P14 was lower than that at other ages (**Figure 1 A and B**). Thus, in the following analyses, we focused on the older ages, namely, P28, P42, P56, and P365. We found that the density of PV+ cells was higher in the OFC than that in the PrL at both P28 and P365, and the soma area of PV+ cells in the OFC was larger than that in the PrL at P28 (**Figure 1C**). These results indicate different age-related changes in PV+ cells in the two regions.

**Figure 1.**
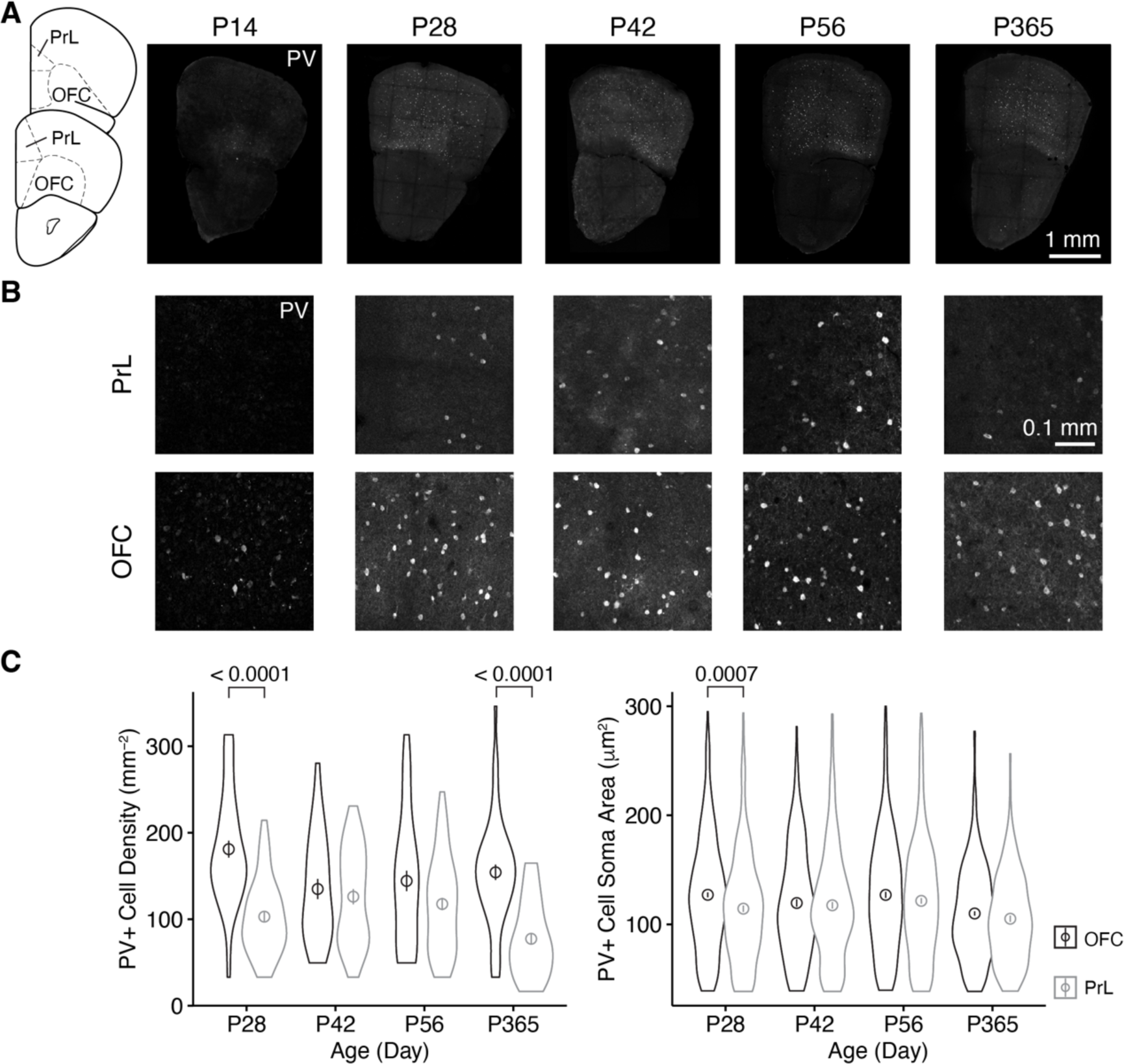
The distribution of PV+ neurons in the OFC and PrL at different ages of mice. **A**, Brain sections containing the OFC and PrL with PV+ neurons immunolabeled with anti-Parvalbumin at different ages. Inset at left, atlas indicating locations of the brain sections. Scale bar, 1 mm. **B**, Blow-up of the OFC and PrL regions at different ages. Scale bar, 0.1 mm. **C**, Comparisons of the cell density and soma area of PV+ neurons in the OFC and PrL regions (Left, PV+ cell density: Two-way ANOVA, *F*(region) = 40.4, *p* = 6.84 × 10^-10^, *F*(age) = 4.5, *p* = 0.004, *F*(region: age) = 7.2, *p* = 0.0001; *post hoc* Tukey’s test: P28 of OFC vs P28 of PrL, *p* = 3.95 × 10^-9^; P42 of OFC vs P42 of PrL, *p* = 0.55; P56 of OFC vs P56 of PrL, *p* =0.06; P365 of OFC vs P365 of PrL, *p* = 1.75 × 10^-9^. Right, PV+ cell soma area: Two-way ANOVA, *F*(region) = 12.0, *p* = 5.52 × 10^-4^, *F*(group) = 11.8, *p* = 1.14 × 10^-7^, *F* (region: group) = 1.3, *p* = 0.26; *post hoc* Tukey’s test: P28 of OFC vs P28 of PrL, *p* =7.36 × 10^-4^; P42 of OFC vs P42 of PrL, *p* = 0.65; P56 of OFC vs P56 of PrL, *p* = 0.15; P365 of OFC vs P365 of PrL, *p* = 0.18. P28 of OFC: cell number, n = 528; section, n = 44; P28 of PrL: cell number, n = 279; section, n = 45; P42 of OFC: cell number, n = 220; section, n = 27; P42 of PrL: cell number, n = 281; n = 37; P56 of OFC: cell number, n = 343; section, n = 40; P56 of PrL: cell number, n = 426; section, n = 60; P365 of OFC: cell number, n = 449; section, n = 48; P365 of PrL: cell number, n = 182; section, n = 39; N = 3/group). Circles and bars in violin plots denote the mean ± sem.

### 3.2 The age-related changes of dopamine D1 and D2 receptor expressions in PV+ neurons in the OFC and PrL

We examined D1R and D2R expressions in PV+ cells in the OFC and PrL (**Figure 2 – 5**). In the OFC, both the density and size of D1R puncta in PV+ cells increased at P42 and P56 when compared with those of P28 and then receded to exist at P365 (**Figure 2**), the expression of D2R showed a similar age-related pattern, but PV+ cells at P365 still showed D2R expression (**Figure 3**).

**Figure 2.**
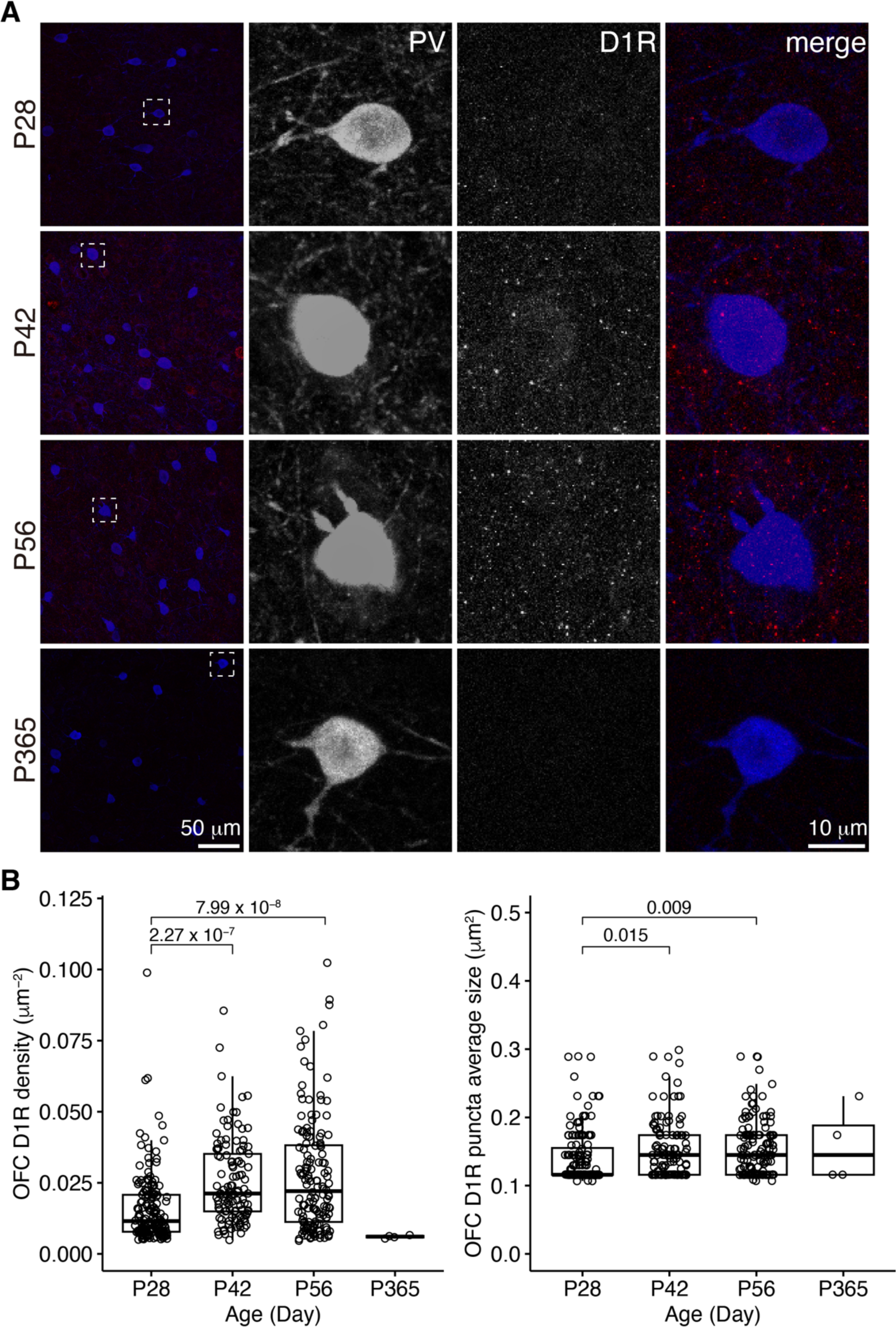
The expression of D1R in PV+ neurons of the OFC. **A**, The characteristic expression of D1R in PV+ neurons in the OFC at different ages. Three panels on the right show the blow-up of the dashed square in the left panel. Scale bars, 50 and 10 μm. **B**, The density and puncta average size of D1R expressed by PV+ neurons in the OFC (Left, OFC D1R density: Welch one-way ANOVA, *F* = 24.8, *p* = 1.56 × 10^-10^; *post hoc* Tukey’s test: P28 vs P42, *p* = 2.27 × 10^-7^; P28 vs P56, *p* = 7.99 × 10^-8^; P42 vs P56, *p* = 0.71. Right, OFC D1R puncta average size: Welch one-way ANOVA, *F* = 6.0, p = 0.003; *post hoc* Tukey’s test: P28 vs P42, *p* = 0.02; P28 vs P56, *p* = 0.009; P42 vs P56, *p* = 0.99. P28, n = 160; P42, n = 113; P56, n = 142; P365, n = 4; N = 3/group).

**Figure 3.**
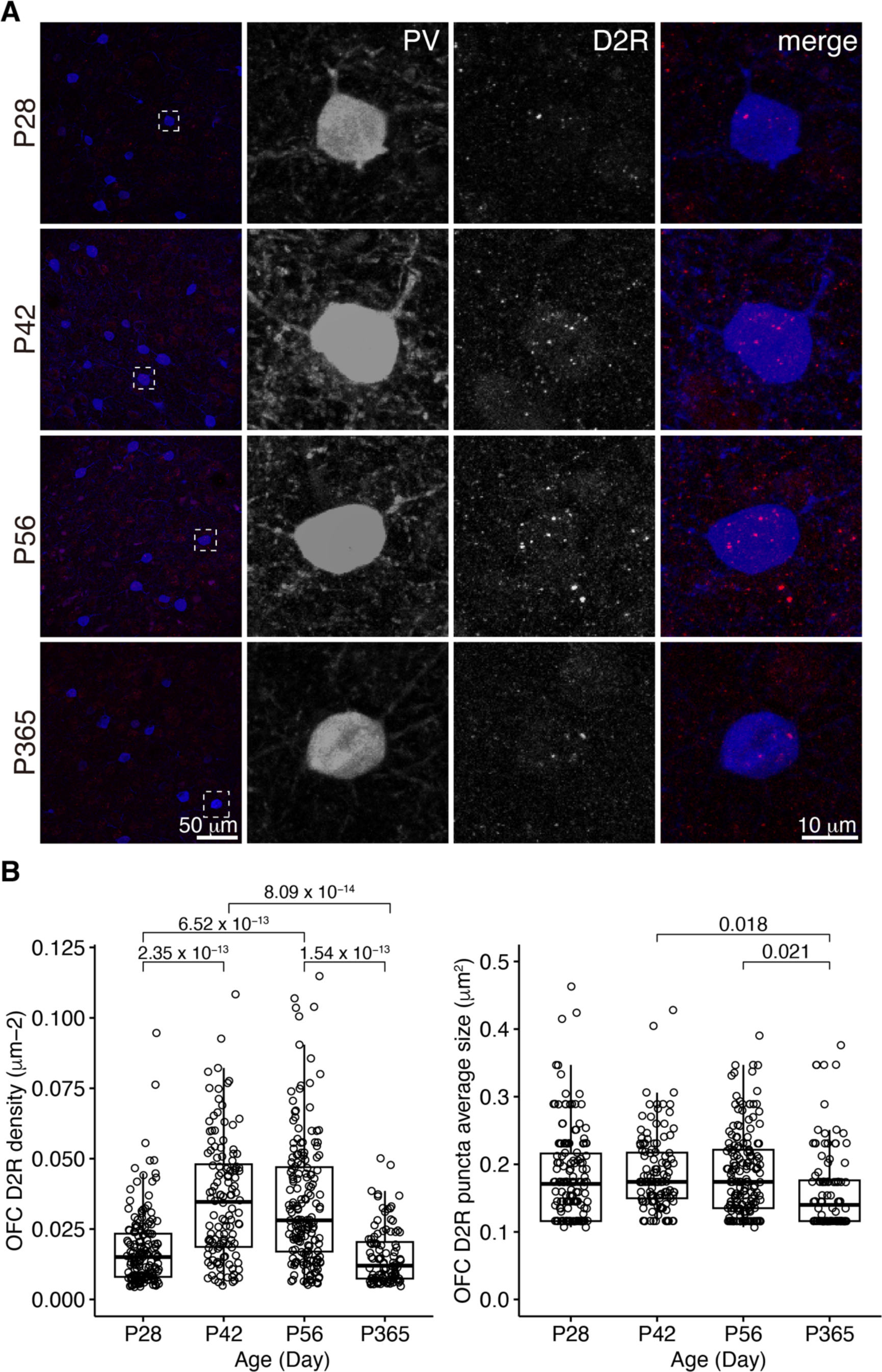
The expression of D2R in PV+ neurons of the OFC. **A**, The expression of D2R in PV+ neurons. Three panels on the right show the blow- up of the dashed square in the left panel. Scale bars, 50 and 10 μm. **B**, The density and puncta average size of D2R expressed by PV+ neurons in the OFC (Left, OFC D2R density: Welch ANOVA, *F* = 51.6, *p* = 8.55 × 10^-27^; *post hoc* Tukey’s test: P28 vs P42, *p* = 2.35 × 10^-13^; P28 vs P56, *p* = 6.52 × 10^-13^; P28 vs P365, *p* = 0.39; P42 vs P56, *p* = 0.96; P42 vs P365, *p* = 8.09 × 10^-14^; P56 vs P365, *p* = 1.54 × 10^-13^. Right, OFC D2R puncta average size: Welch ANOVA, *F* = 3.5, *p* = 0.02; *post hoc* Tukey’s test: P28 vs P42, *p* = 0.83; P28 vs P56, *p* = 0.88; P28vs P365, *p* = 0.15; P42 vs P56, *p* = 1.00; P42 vs P365, *p* = 0.02; P56 vs P365, *p* = 0.02. P28, n = 162; P42, n = 124; P56, n = 171; P365, n = 100; N = 3/group).

In the PrL, the D1R expression in PV+ neurons were scarce across ages (**Figure 4**), but the D2R expression in PV+ neurons showed age-related changes similar to that of the OFC (**Figure 5**).

**Figure 4.**
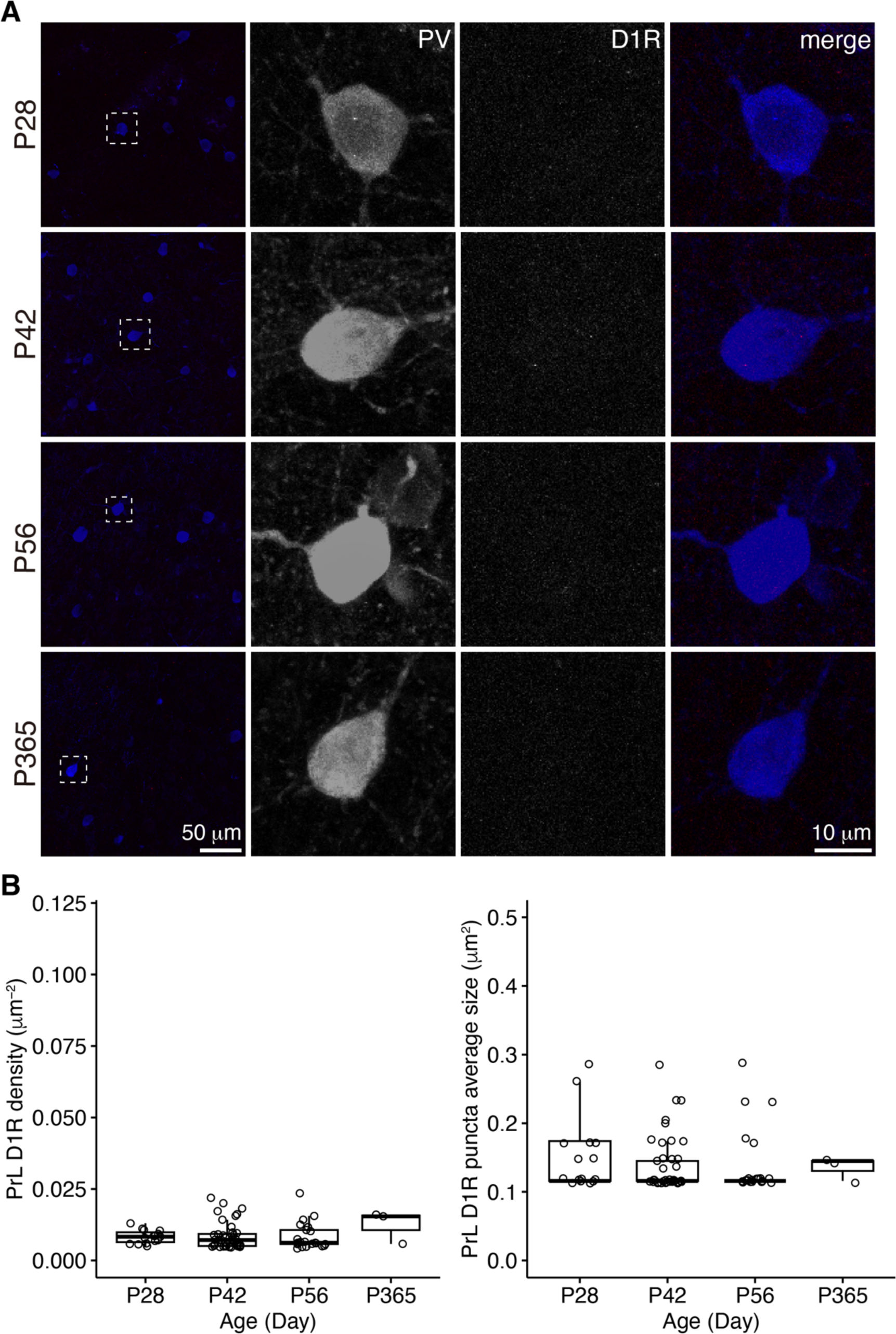
The expression of D1R in PV+ neurons of the PrL. **A**, The expression of D1R in PV+ neurons. Three panels on the right show the blow- up of the dashed square in the left panel. Scale bars, 50 and 10 μm. **B**, The density and puncta average size of D1R expressed by PV+ neurons in the PrL (Left, PrL D1R density: Welch one-way ANOVA, *F* = 0.1, *p* = 0.94. Right, PrL D1R puncta average size: Welch one-way ANOVA, *F* = 0.3, *p* = 0.73. P28, n = 15; P42, n = 40; P56, n = 21; P365, n = 3; N = 3/group).

**Figure 5.**
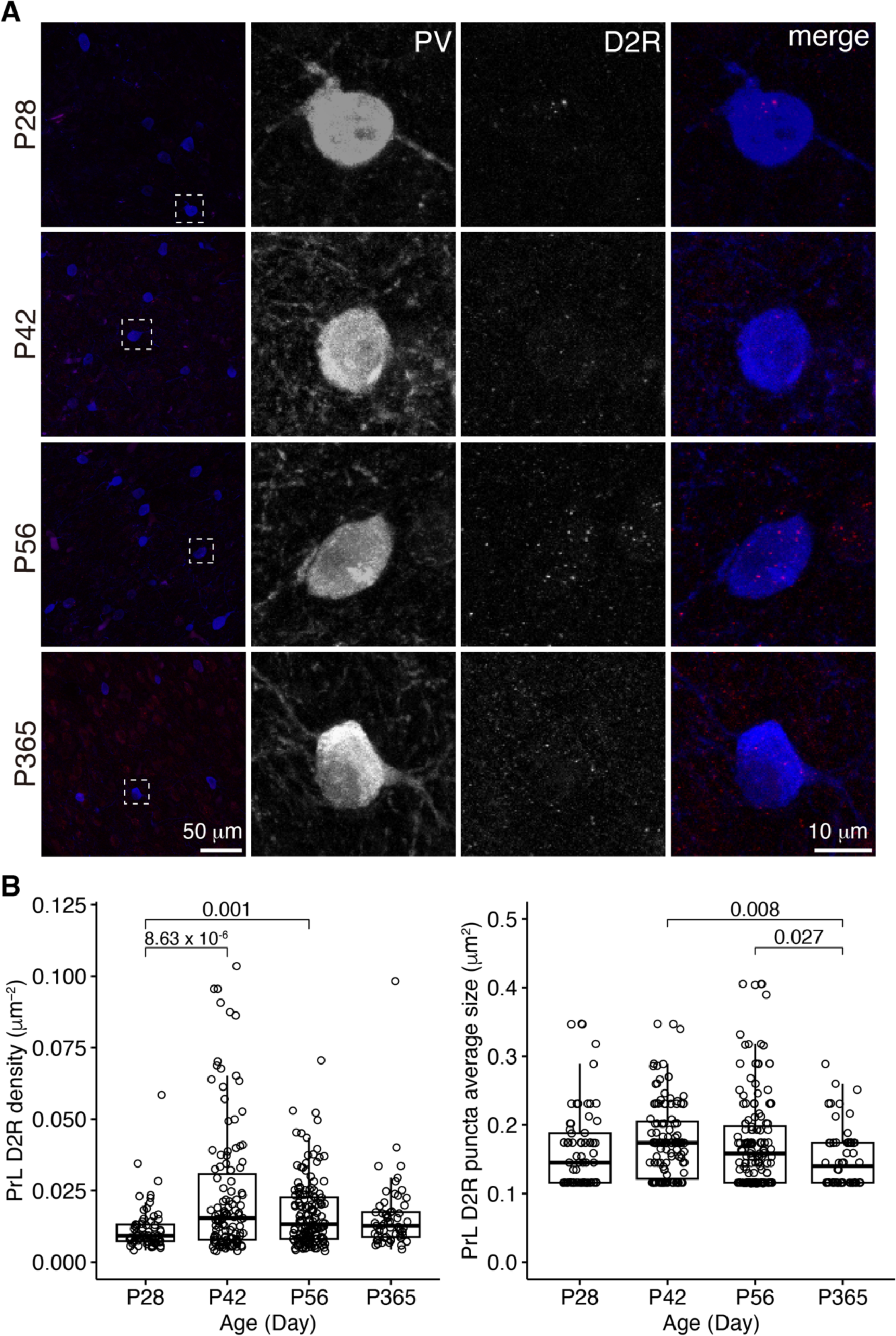
The expression of D2R in PV+ neurons of the PrL. **A**, The expression of D2R in PV+ neurons. Three panels on the right show the blow- up of the dashed square in the left panel. Scale bar, 50 μm, 10 μm. **B**, The density and puncta average size of D2R expressed by PV+ neurons in the PrL(Left, PrL D2R density: Welch one-way ANOVA, *F* = 10.1, *p* = 3.43 × 10^-6^; *post hoc* Tukey’s test: P28 vs P42, *p* = 8.31 × 10^-6^; P28 vs P56, *p* = 0.001; P28 vs P365, *p* = 0.17; P42 vs P56, *p* = 0.03; P42 vs P365, *p* = 0.03; P56 vs P365, *p* = 0.94. Right, PrL D2R puncta average size: Welch one-way ANOVA, *F* = 4.12, *p* = 0.007; *post hoc* Tukey’s test: P28 vs P42, *p* = 0.58; P28 vs P56, *p* = 0.75; P28 vs P365, *p* = 0.48; P42 vs P56, *p* = 0.99; P42 vs P365, *p* = 0.008; P56 vs P365, *p* = 0.03. P28, n = 67; P42, n = 119; P56, n = 148; P365, n = 60; N = 3/group).

### 3.3 Comparisons of D1R and D2R expression in PV+ neurons of the OFC and PrL

We categorized PV+ neurons in the OFC and PrL based on the expression patterns of D1R and D2R into four groups, namely D1R-D2R-, D1R+D2R-, D1R- D2R+, and D1R+D2R+ (**Figure 6A** and **Table 1**). We found that PV+ neurons in these regions showed different expression patterns of the D1R and D2R. Correlation analysis of the density of D1R and D2R in each cell of the PrL and OFC across ages further showed that overall PV+ neurons in the OFC expressed more D1R than D2R, while PV+ neurons in the PrL showed the opposite pattern (**Figure 6B**). Further analyses at different ages confirmed a similar region-specific pattern except for P365 when the expression D1R in PV+ neurons in both brain regions receded (**Figure 6 C** and **D**).

**Figure 6.**
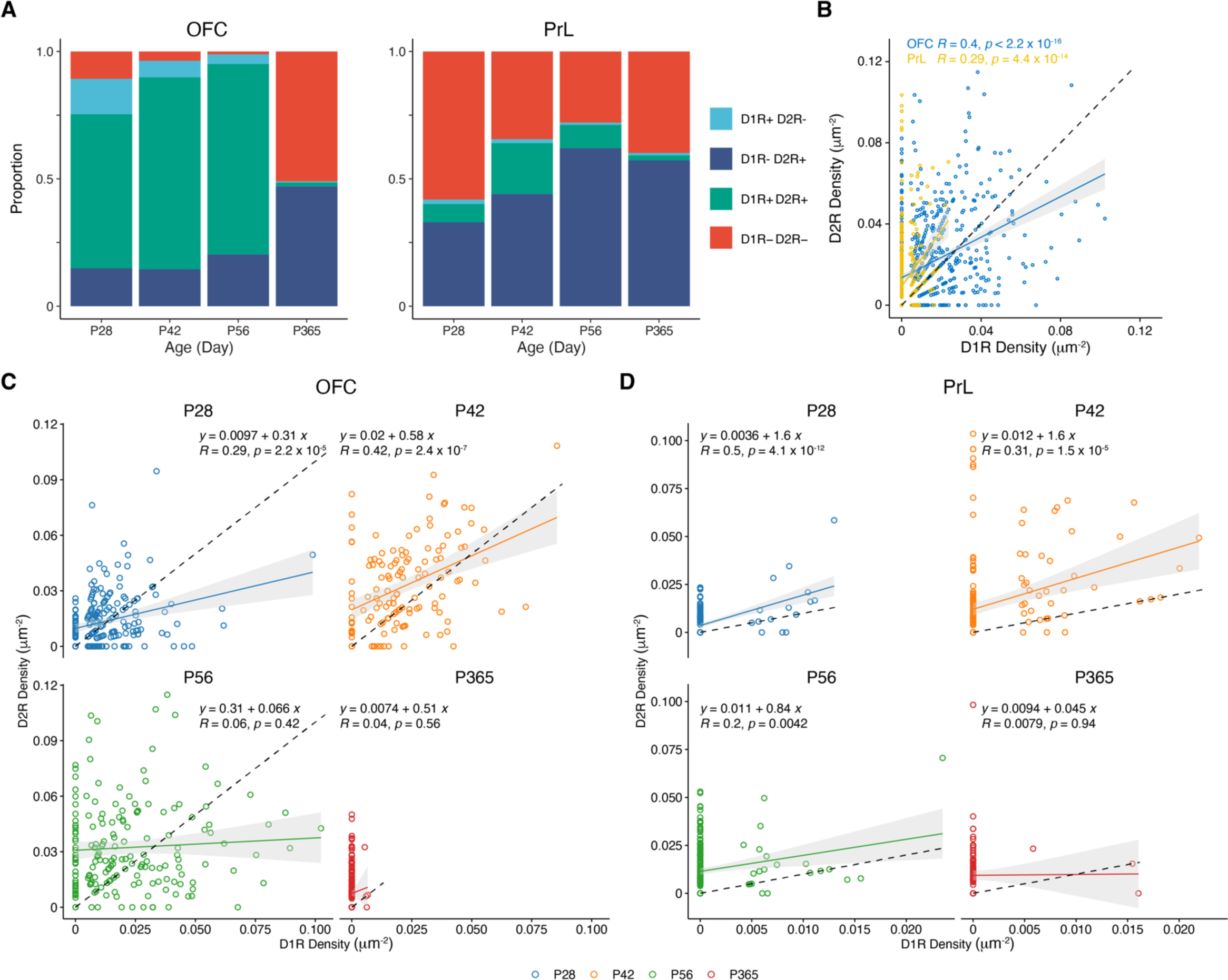
Correlation analysis of the expressions of D1R and D2R in PV+ neurons. **A**, The proportion of D1R and D2R expressed by PV+ neurons in the OFC and PrL (N = 3/group). **B**-**D**. Correlation analysis of density of D1R and D2R expressions in PV+ neurons in the OFC and PrL (**B**, OFC, n = 741; PrL, n = 663. **C**, P28, n = 215; P42, n = 138; P56, n = 182; P365, n = 206. **D**, P28, n = 167; P42, n = 186; P56, n = 208; P365, n = 102; N = 3/group).

**Table 1.**
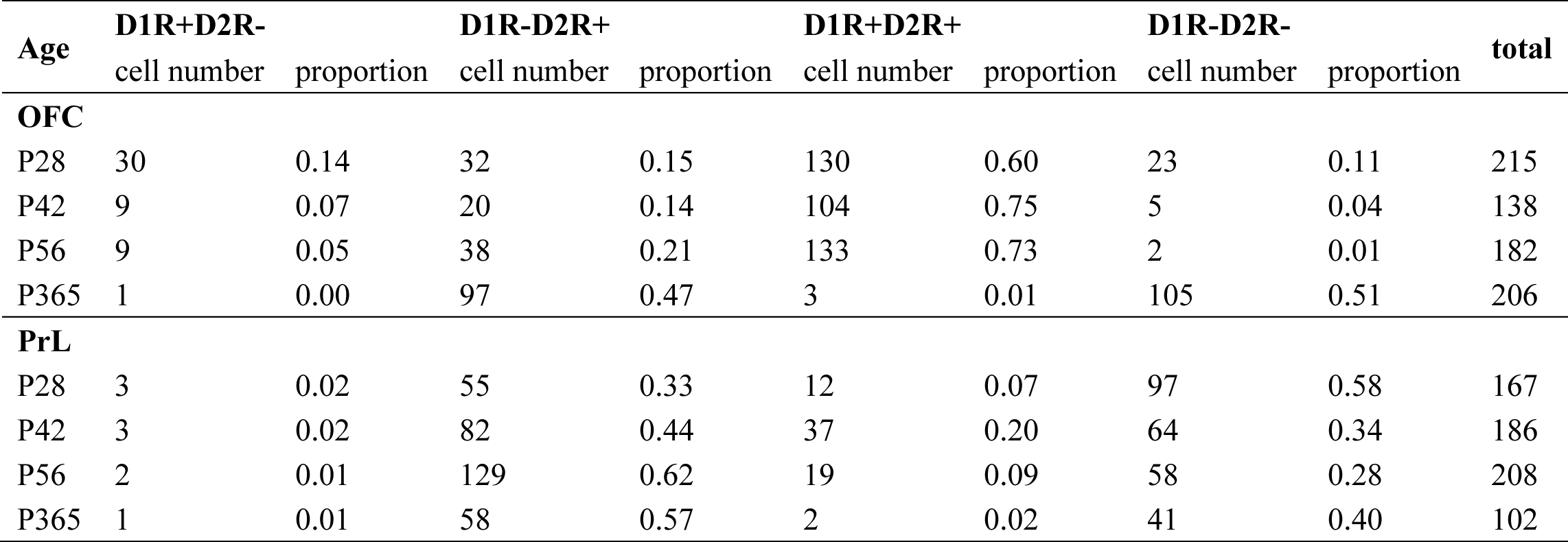
Summary of cell numbers of OFC and PrL.

We then compared the expression of D1R and D2R in PV+ cells of the four groups in the OFC and the PrL (**Figure 7**). Of the four groups, only D1R-D2R+ and D1R+D2R+ groups showed differences between the two brain regions (**Figure 7B** and **C**). Of the D1R+D2R+ group, the density of D1R in PV+ cells was higher in the OFC than that in the PrL at P42 and P56 (**Figure 7C**), while the density of D2R in PV+ cells was higher in the OFC than that in the PrL at P56 which is similar to the density of D2R of D1R-D2R+ group cells (**Figure 7B**).

**Figure 7.**
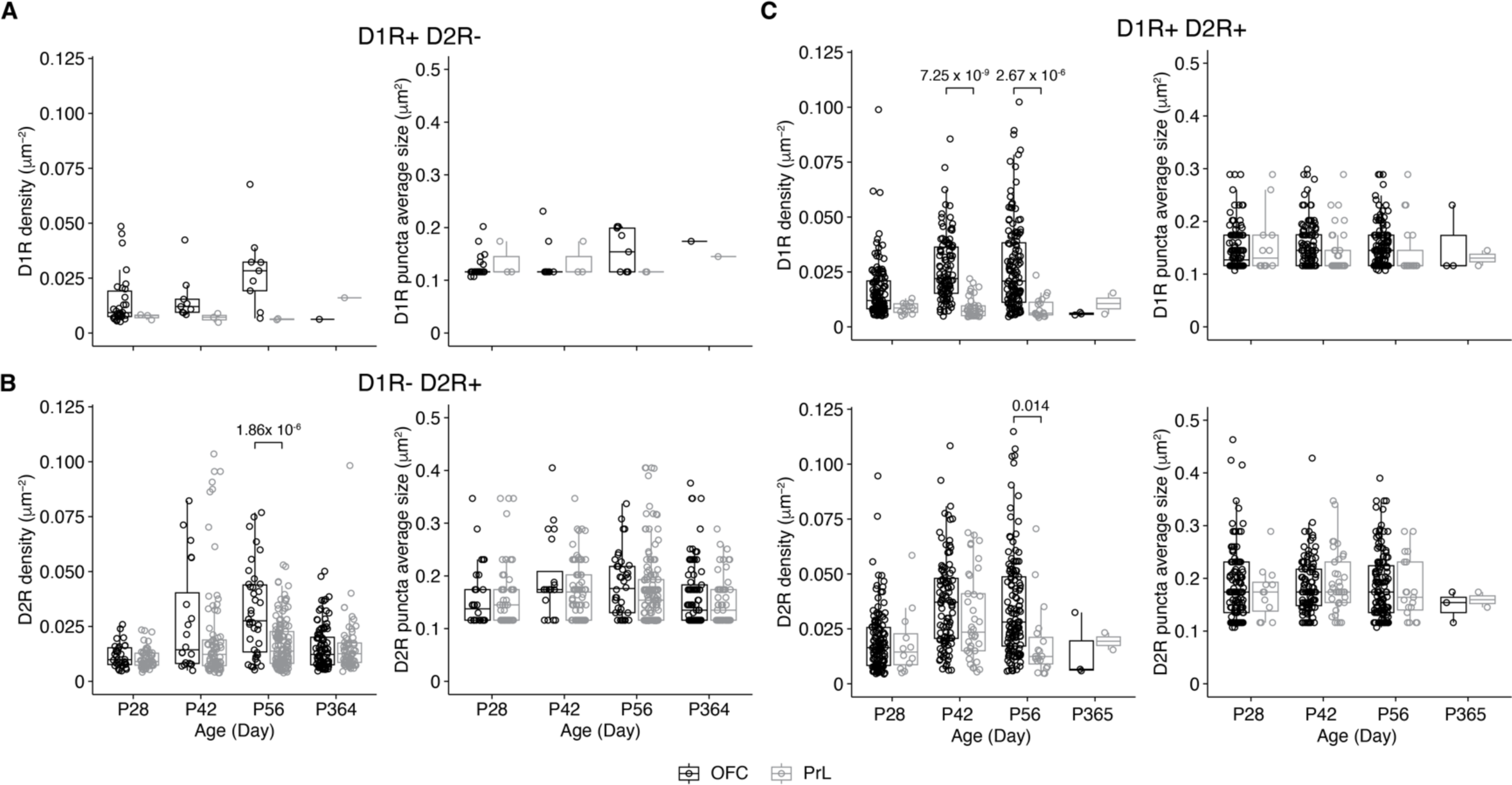
Comparisons of D1R and D2R expressions of PV+ neurons of the PrL and OFC. **A**, Comparative analysis of the density of PV+ neurons expressing only D1R in the OFC and PrL (Left, D1R density: Two-way ANOVA, *F*(region) = 0.9, *p* = 0.34, F (Age) = 3.3, *p* = 0.03, *F*(region : Age) = 1.0, *p* = 0.39; *post hoc* Tukey’s test: OFC vs PrL, *p* = 0.04; P28 vs P56, *p* = 0.05. Right, D1R puncta average size: Two-way ANOVA, *F*(region) = 0.5, *p* = 0.50, *F*(Age) = 3.6, *p* = 0.02, *F*(region : Age) = 1.2, *p* = 0.31. OFC: P28, n = 30; P42, n = 9; P56, n = 9; P365, n = 1; PrL: P28, n = 3; P42, n = 3; P56, n = 2; P365, n = 1; N = 3/group). **B**, Comparative analysis of the density of PV+ neurons expressing only D2R in the OFC and PrL (Left, D2R density: Two-way ANOVA, *F*(region) = 0.2, *p* = 0.69, *F*(Age) = 16.1, *p* = 5.13 × 10^-10^, *F*(region : Age) = 6.5, *p* = 2.46 × 10^-4^; *post hoc* Tukey’s test: OFC vs PrL, *p* = 0.03; P28 vs P42, *p* = 7.47 × 10^-7^; P28 vs P56, *p* = 3.91 × 10^-6^; P42 vs P365, *p* = 4.24 × 10^-4^; P56 vs P365, *p* = 2.58 × 10^-3^; P28 of OFC vs P42 of OFC, *p* = 5.36 × 10^-3^; P28 of OFC vs P56 of OFC, *p* = 6.09×10^-7^; P28 of PrL vs P42 of OFC, *p* = 3.49 × 10^-4^; P28 of PrL vs P42 of PrL, *p* = 0.002; P28 of PrL vs P56 of OFC, *p* = 6.25×10^-10^; P42 of OFC vs P365 of OFC, *p* = 0.02; P42 of PrL vs P56 of OFC, *p* = 0.003; P56 of OFC vs P56 of PrL, *p* = 1.86 × 10^-6^; P56 of OFC vs P365 of OFC, *p* = 3.46 × 10^-7^; P56 of OFC vs P365 of PrL, *p* = 9.18 × 10^-6^. Right, D2R puncta average size: Two-way ANOVA, *F*(region) = 0.3, *p* = 0.61, *F*(Age) = 2.7, *p* = 0.05, *F*(region: Age) = 0.9, *p* = 0.43; *post hoc* Tukey’s test: P56 vs P365, *p* = 0.04. OFC: P28, n = 32; P42, n = 20; P56, n = 38; P365, n = 97; PrL: P28, n = 55; P42, n = 82; P56, n = 129; P365, n = 58; N = 3/group). **C**, Comparative analysis of the density of D1R and D2R co-expressed by PV+ neurons in OFC and PrL (Top-left, D1R density: Two-way ANOVA, *F*(region) = 2.9, *p* = 0.09, *F*(Age) = 14.8, *p* = 3.68 × 10^-9^, *F*(region: Age) = 2.04, *p* = 0.11; *post hoc* Tukey’s test: OFC vs PrL, *p* < 0.00001; P28 vs P42, *p* = 1.44 × 10^-5^; P28 vs P56, *p* = 3.33 × 10^-7^; P28 of OFC vs P42 of OFC, *p* = 1.9 × 10^-5^; P28 of OFC vs P56 of OFC, *p* = 3.48×10^-7^; P28 of PrL vs P42 of PrL, *p* =0.003; P28 of PrL vs P56 of OFC, *p* = 0.001; P42 of OFC vs P42 of PrL, *p* = 7.25 × 10^-8^; P42 of OFC vs P56 of PrL, *p* = 1.08 × 10^-4^; P42 of PrL vs P56 of OFC, *p* = 4.53 × 10^-9^; P56 of OFC vs P56 of PrL, *p* = 2.67 × 10^-5^. Top-right, D1R puncta average size: Two-way ANOVA, *F*(region) = 0.90, *p* = 0.34, *F*(Age) = 1.8, *p* = 0.15, *F*(region: Age) = 1.2, *p* = 0.30. Bottom-left, D2R density: Two-way ANOVA, *F*(region) = 0.005, *p* = 0.95, *F*(Age) = 21.4, *p* = 6.05 × 10^-13^, *F*(region: Age) = 1.8, *p* = 0.15; *post hoc* Tukey’s test: OFC vs PrL, *p* = 0.04; P28 vs P42, *p* < 0.00001; P28 vs P56, *p* = 1.27 × 10^-8^; P28 of OFC vs P42 of OFC, *p* = 2.46 × 10^-10^; P28 of OFC vs P56 of OFC, *p* = 4.71×10^-9^; P28 of PrL vs P42 of OFC, *p* = 0.04; P42 of OFC vs P56 of PrL, *p* = 0.003; P56 of OFC vs P56 of PrL, *p* = 0.01. Bottom-right, D2R puncta average size: Two-way ANOVA, *F*(region) = 0.5, *p* = 0.50, *F*(Age) = 0.40, *p* = 0.76, *F*(region: Age) = 0.4, *p* = 0.74. OFC: P28, n = 130; P42, n = 104; P56, n = 133; P365, n = 3; PrL: P28, n = 12; P42, n = 37; P56, n = 19; P365, n = 2; N = 3/group).

## 4 Discussion

In the present study, we showed that PV+ neurons in the OFC expressed both D1 (D1R) and D2 (D2R) receptors, while those in the PrL mainly expressed D2 receptors. D1 and D2 receptors are known for their different roles in regulating the excitability and GABAergic transmission of interneurons in the PFC. Studies suggest that D1- and D2-like dopamine receptors regulate interneuron activity and GABAergic transmissions through different mechanisms. For example, D1R but not D2R agonist enhanced spontaneous IPSCs (sIPSCs), while D2R agonist reduced miniature IPSCs (Li, Lin, Yang, & Xie, 2015; Seamans et al., 2001). These data suggest that D1-like dopamine receptors increased the excitability of interneurons, and consistent with this finding, blockage of D1-like but not D2-like dopamine receptors abolished dopamine-induced increased excitability of fast-spiking interneurons (Gorelova, Seamans, & Yang, 2002). On the other hand, a study showed that the activation of D1-like dopamine receptors inhibited evoked-GABAergic transmission, while agonists of D2- like dopamine receptors had no effect (Gonzalez-Islas & Hablitz, 2001). Considering the different dopamine receptor expression patterns of PV+ neurons in the PrL and the OFC, dopamine might induce different functional changes of the PV+ neurons in the PRL and the OFC, which would contribute to different changes of network activity in these brain regions.

In the brain, D1R and D2R can form heterodimer (Perreault, Hasbi, O’Dowd, & George, 2014), and studies suggest the activation of such heterodimer recruits Gαq/11 and releases calcium from the internal stores (Hasbi et al., 2009; Rashid et al., 2007). Interestingly, dopamine increased the excitability of layer I interneurons, which was mimicked with the co-application of D1-like and D2-like dopamine receptor agonists (Wu & Hablitz, 2005), indicating a possible synergic effect of D1- and D2-like receptor interaction. In the present study, we also observed higher proportions of PV+ neurons with D1 and D2 receptor co-expressions in the OFC than those in the PrL; however, whether these neurons also showed a similar synergic effect upon dopamine activation needs further investigation. Furthermore, whether these neurons expressed D1/D2 receptor heteromers is unclear. D1/D2 receptor heteromers have been reported in the prefrontal cortex (Hasbi et al., 2020) ; nevertheless, whether D1/D2 receptor heteromers are present *in vivo* is still under debate (Frederick et al., 2015).

The PRL and OFC are two brain regions that regulate emotions, decision-making, and other cognitive processes. While they share some similarities, there are also some key differences between these two regions. The OFC is suggested as the first stage of cortical processing of the reward value-related information, with neurons that respond to the outcome and the expected value (Padoa-Schioppa & Conen, 2017; Rolls, 2019, 2021). It is worth noting that those expected-value neurons do not reflect prediction error as they keep responding to the expected reward without prediction error.

Furthermore, studies have shown that OFC is involved in decision-making by representing rewards, punishes, and errors during decision-making (Rolls, 2019), while the PrL is more related to prediction error (Casado-Roman, Carbajal, Perez- Gonzalez, & Malmierca, 2020). The higher proportions of D1R+ PV+ neurons in the OFC might contribute to these processes.

We also observed different development changes of dopamine receptors in the PV+ neurons between the OFC and PrL. More than 80 % of PV+ neurons in the OFC expressed dopamine receptors at P28, while less than 50 % of those in the PrL did. At the following ages of P42 and P56, while the percentages of dopamine receptor-expressing PV+ neurons increased in both the PrL and OFC, the patterns of change differed between the two brain regions. In the OFC, the percentage of D1R+ D2R+ PV+ neurons increased with age. Meanwhile, the percentage of D1R-D2R+ PV+ neurons increased in the PrL. For mice at P365, PV+ neurons in both the PrL and OFC showed a significant reduction in the expression of dopamine receptors, and the majority of those neurons only expressed D2R. Studies examined D1R and D2R expression in the cerebral cortex also found D2R expression did not change from birth to 70 weeks-old while D1R expression reached a plateau around 2 – 3 weeks-old then decreased in the prefrontal cortex in rodents (Leslie, Robertson, Cutler, & Bennett, 1991; Rani & Kanungo, 2006). While these studies analyzed the overall expression of those two types of dopamine receptors, the present study focused on the expression of these receptors on PV+ neurons. Considering that ∼ 30% of inhibitory neurons in the cortex are PV+ neurons and ∼ 20% of cortical neurons are inhibitory, the PV+ neurons might exhibit a different expression pattern than the overall expressions of D1R and D2R in the cortex.

In the present study, we showed the age- and region-specific expression patterns of D1 and D2 dopamine receptors in PV+ neurons of the orbitofrontal and prelimbic regions of the prefrontal cortex. Our results showed that while D1R+ PV+ neurons in both regions decreased along with aging, PV+ neurons in the OFC showed higher density and percentage of D1R and D2R expression than that of the PrL through adulthood. These results provide anatomical evidence for the understanding PV+ neuron functions in these regions in reward-related brain functions.

## Acknowledgments

This research was supported by Ministry of Science and Technology of China (2019YFA0706201, WZ) and National Natural Science Foundation of China (32170960, WZ).

## Author Contributions

All authors had full access to all the data in the study and take responsibility for the integrity of the data and the accuracy of the data analysis. Study concept and design: WZ; Acquisition of data: JD, XW, ZH, JT; Analysis and interpretation of data: JD, WZ; Drafting of the manuscript: WZ; Critical revision of article for important intellectual content: JD; Obtained funding: WZ; Study supervision: WZ.

